# Modeling the biomechanics of cells on microcarriers in a stirred-tank bioreactor

**DOI:** 10.1101/2022.08.31.505282

**Authors:** Jaro Camphuijsen, Fernando J. Cantarero Rivera, Greg Potter, Chris Clark, Jiajia Chen, Simon Kahan, Boris Aguilar

## Abstract

Highly productive and efficient growth of biomass in bioreactors is an essential bioprocess outcome in many industrial applications. In the nascent cultivated meat industry, large-scale biomass creation will be critical given the size of demand in the conventional meat and seafood sectors. However, there are many challenges that must be overcome before cultivated meat and seafood become commercially viable including cost reductions of cell culture media, bioprocess design innovation and optimization, and scaling up in the longer term. Computational modelling and simulation can help to address many of these challenges, and can be a far cheaper and faster alternative to performing physical experiments. Computer modelling can also help researchers pinpoint system interactions that matter, and guide researchers to identify those parameters that should be changed in later designs for eventual optimization. In this work, a computational model that combines agent-based modeling and computational fluid dynamics was developed to study biomass growth as a function of the operative conditions of stirred-tank bioreactors. The focus was to analyze how the mechanical stress induced by rotor speed can influence the growth of cells attached to spherical microcarriers. The computer simulation results reproduced observations from physical experiments that high rotor speeds reduce cell growth rates and induce cell death under the high mechanical stresses induced at these stir speeds. Moreover, the results suggest that modeling both cell death and cell quiescence are required to recapitulate these observations from physical experiments. These simulation outcomes are the first step towards more comprehensive models that, in combination with experimental observations, will improve our knowledge of biomass production in bioreactors for cultivated meat and other industries.

## Introduction

Increasing worldwide demand for meat is driving the growth of environmentally detrimental factory farming, aquaculture, and mass capture of fish and other marine life [1]. Cultivated meat and seafood is a potentially more sustainable alternative to factory farming and fisheries practices that could mitigate land use, disease spread, greenhouse gas emissions, and water contamination [2,3]. As a nascent technology, however, several challenges must be overcome before cultivated meat is available worldwide. One of the most fundamental challenges in commercializing cell-based meat is producing biomass at a cost lower than that of animal husbandry and slaughter [2]. At present, both the cost of cell culture media and industrial-scale perfusion bioreactors production make cultivated meat orders of magnitude more expensive than the conventional alternatives [4]. To be cost-competitive, cheaper media formulations of non-animal origin (serum-free) will be a necessity. Similarly, more economical bioreactor designs and configurations must be developed to overcome the cost hurdle [4]. Unfortunately, there are currently no commercial cultivated meat bioreactors or bioprocesses in existence [5]. Cultivated meat bioreactor designs do have parallels in cell therapy and vaccine production bioprocesses, but the cultivated meat use cases are distinctive because of the far greater number of cells that must be produced to create even one serving of meat compared to delivering a dose of therapy [6].

The starter cells are a primary area of focus for cultivated meat bioreactor design and prototyping. These cells can be sensitive to turbulent flow and the resulting mechanical forces that induce detachment, damage, apoptosis, and untimely differentiation [7]. Therefore, bioreactor design considerations must accommodate starter cell mechanical force sensitivities while still achieving sufficient mass transfer, oxygen transfer, and adequate CO_2_ removal [6,8]. Maintaining a favorable micro and macro-environment for cells without subjecting them to excessive mechanical stress due to high stirring speed will require innovation in, and optimization of, bioreactor designs and processes.

Virtual prototyping and experimentation through computer simulations promise to accelerate and lower the cost of progress [9]. Virtual experiments that replicate actual bioreactor and biological behaviors are not currently possible. First, it is necessary to develop predictive models of the bioreactor environment [9]. The main challenge in developing a predictive model of cells growing in a bioreactor is the complexity of the bioreactor environment and cell behavior. Media flow dynamics, forces, and mixing of media components need to be incorporated into the model. Simultaneously, growing cells that consume nutrients, excrete waste, proliferate, and die introduce an additional layer of complexity [9]. The high density of cells needed for efficient production of meat comes with more interactions between cells in the bioreactor and potential aggregation of cells in suspension and microcarrier cultures. Thus, despite prior work using computer modeling and simulation to predict bioreactor behavior for numerous designs and configurations [9], an altogether new multiscale methodology accounting for phenomena happening at diverse spatial and temporal scales seems necessary to be able to include these cell-cell interactions.

Two well-established computational modeling approaches show considerable promise as a means to encapsulate the complexity of the bioreactor environment and cell behavior - computational fluid dynamics (CFD) and agent-based modeling (ABM). In cultivated meat production, CFD can be used to simulate and understand the fluid flows in the bioreactors governed by the Navier-Stokes equations [10]. In biological applications, ABM can be used to track the fate of each individual cell by following rules that determine the conditions under which the cell, for example, grows, moves, adheres, divides, differentiates, and dies as a function of the biochemical and physical environment [11].

Stirred-tank reactors are very well-studied, and are being pursued in cultivated meat production [6]. Adherent cells specifically can be grown in stirred-tank systems with microcarriers, but microcarrier-bound cells become sensitive to agitation damage by mechanical forces and small intense eddies [12]. In this study, we present the first co-application of CFD and ABM as a multiscale modeling methodology for cultivated meat bioreactor and bioprocess design by focusing on a simple configuration of a stirred tank bioreactor system [12]. The research paper of Croughan *et al*. [12] provides both empirical and theoretical analyses of hydrodynamic effects on microcarrier cultures of FS-4 cells in a stir-rod bioreactor. In particular, the paper reports distinct trends for the influence of stir speed on biomass accumulation, and thus, growth rate and death rate. Our novel modeling methodology represents a simple framework for the concurrent application of CFD and ABM to simulate the biophysical phenomenon of cell response to increasing bioreactor stir speeds. The framework is constructed in a way that enables expansion to include additional knowledge and should be applicable to other bioreactor and bioprocess designs in cultivated meat and allied industries.

## Materials and Methods

### Bioreactor Setup

In the simulations, the microcarrier cultures were grown in a simple bioreactor set-up that was agitated but was not aerated and had no pH control technology (Fig 1). A cylindrical rod immersed to two-thirds the depth of the liquid and rotated at a constant speed agitates the fluid. Rotational speed was the only parameter varied in the experiments reported by Croughan *et al* [12]. This and other parameters of the bioreactor system are specified (S1 Table). In our model, we needed additional quantities (e.g. microcarrier and cell dimensions) not provided by Croughan *et al* [12], attributed to other sources (S1 Table).

**Fig 1.**
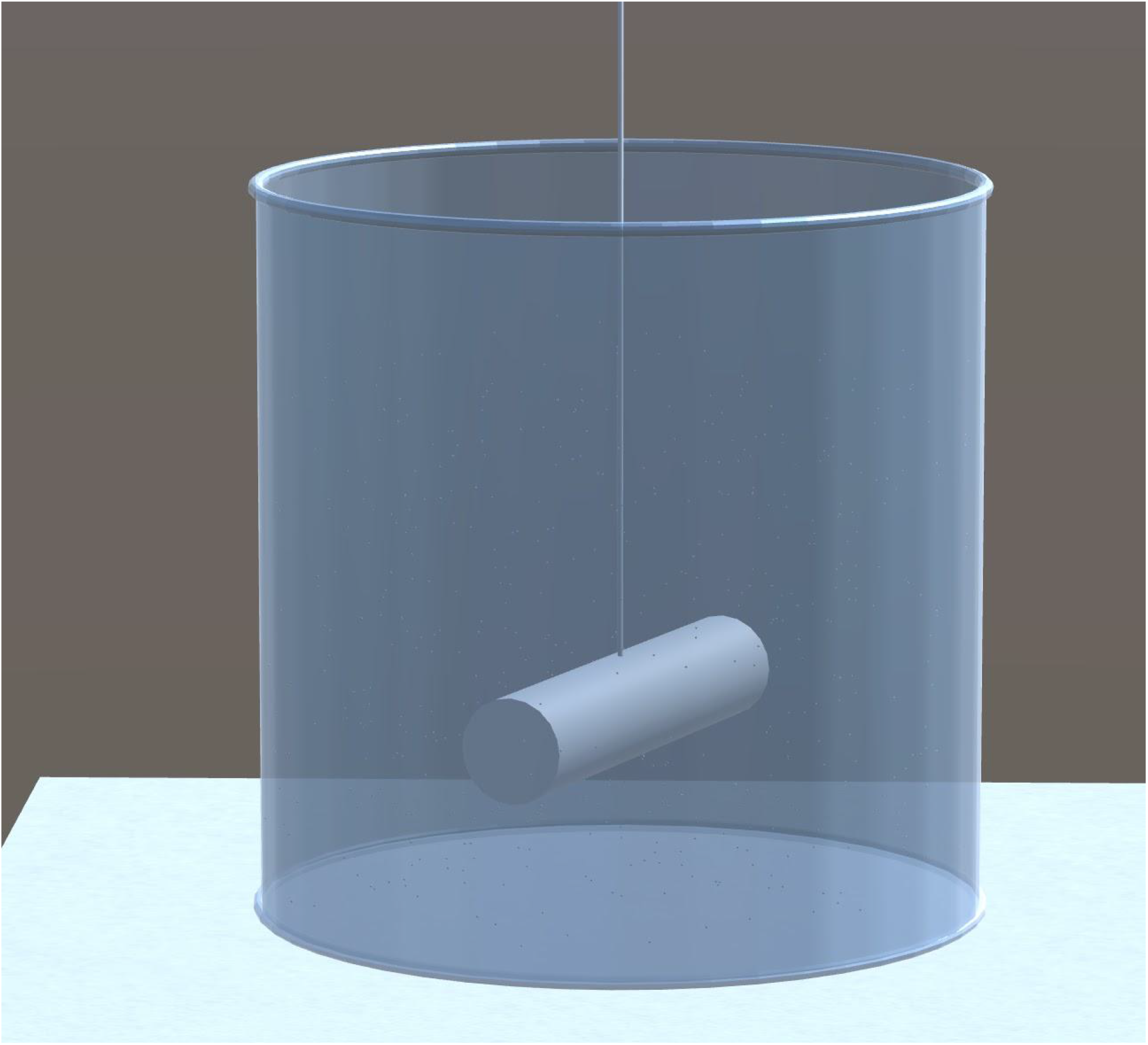
Impression of the bioreactor. Used as boundary conditions in ABM and CFD simulations

Croughan *et al*. [12] studied how growth rate changes as a function of excessive agitation speed. Prior experimental results had established roughly 60 rpm to be a rate sufficient to provide the needed supplies of nutrients and oxygen while higher rates reduced productivity. Croughan *et al*. [12] were focused on rates of 60 rpm and above. They proposed an analytical model in which growth and death rates are exponential over time, resulting in population graphs that are piecewise linear when on a log-scale as shown in Fig 2. In their model, the growth (proliferation) rate exponent of cells is assumed constant throughout all experiments and the death rate is assumed to change with agitation speed. The rates are calculated based on these assumptions to best-fit data sampled from experiments.

**Fig 2.**
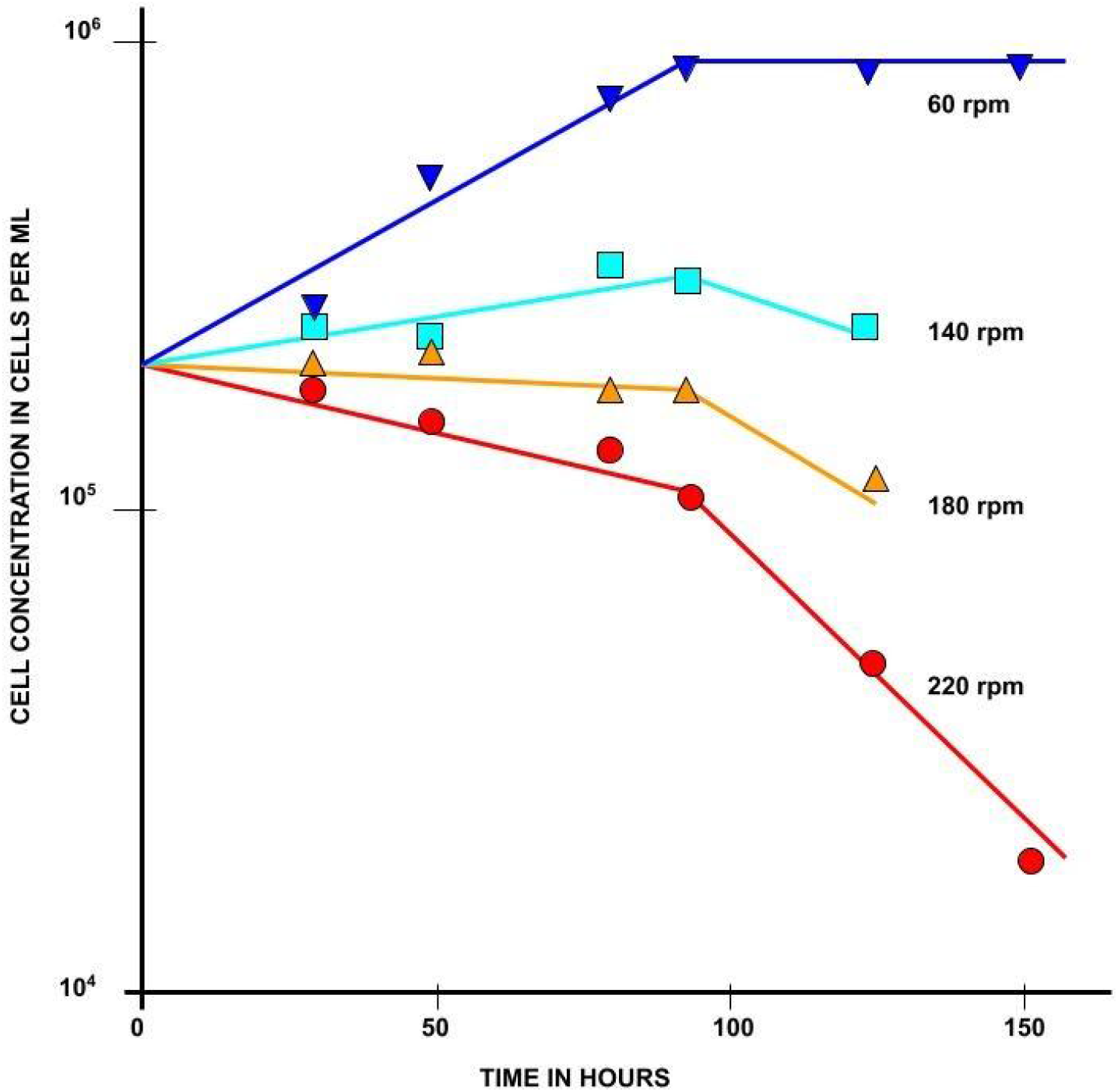
Growth of FS-4 cells on microcarriers at various stirring speeds. All cultures contained 3 g/L Cytodex 1 microcarriers and were in identical 250 ml vessels. This figure is a reproduction of Figure 1 from Croughan et al. [12] and illustrates the set of experiments the simulations in this research are based on.

### Computational Fluid Dynamics (CFD)

#### Geometry

The geometry of the CFD model included a 100 mL cylindrical bioreactor and a cylindrical stirring bar [12], as shown in Fig 3. The cylindrical bioreactor had an internal diameter of 5.5 cm, and the rod impeller had a length of 3.8 cm and a diameter of 0.8 cm. The rod impeller was suspended at one-third of the liquid height. In order to simulate the rotation of the rod impeller, the moving mesh technique was used and an imaginary domain was created within the bioreactor.

**Fig 3.**
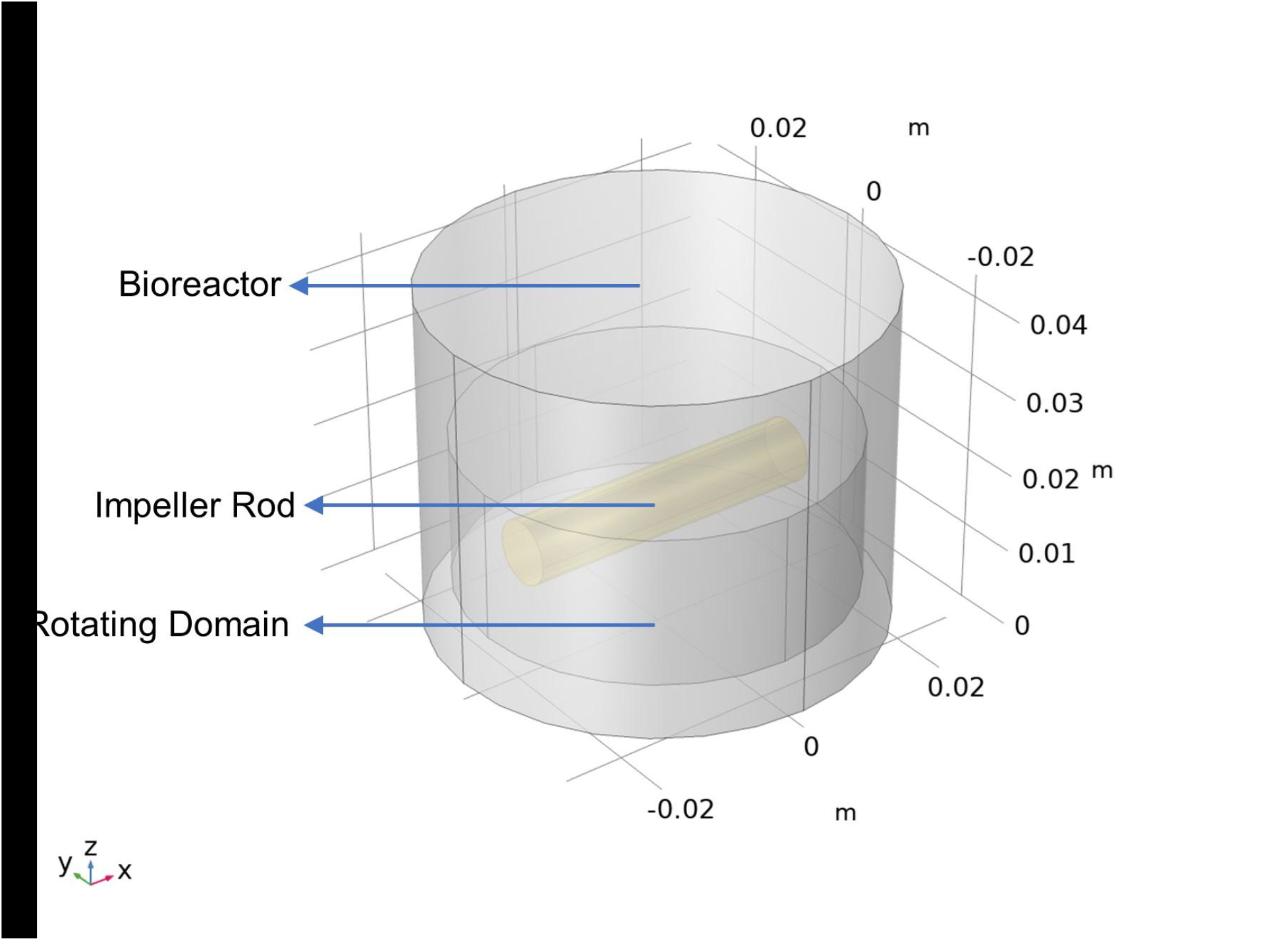
Geometry of the bioreactor with a rod impeller.

#### Governing equations and boundary conditions

The fluid flow in the bioreactor was assumed to be turbulent and a Reynolds-averaged Navier-Stokes (RANS) simulation approach with the standard *k-*ε turbulence model was used to simulate the fluid movement, where *k* represents the turbulent kinetic energy, and *ε* represents the dissipation of turbulent kinetic energy. Since the mixing process is continuous during the whole cell growth period (∼days), the fluid flow reaches a steady-state condition. Therefore, the equations for mass conservation, Newton’s second law, and energy conservation, along with the constitutive laws relating the stress tensor in fluid to the rate of the deformation tensor were used to describe the fluid flow process [10], shown as a general form :

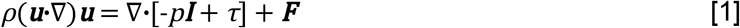

where *ρ* is the density, ***u*** is the velocity vector, *p* is pressure, **I** is the identity matrix, *τ* is the viscous stress tensor, and ***F*** is the volume force vector.

A no slip boundary condition was applied to the bioreactor walls. This wall condition must be included to compensate for the limitations of the *k-*ε turbulence model close to the walls.

#### Material Properties

The material properties of the growth media were assumed to be that of liquid water (ρ=1000 kg.m^−3^, μ=0.001003 Pa.s) and did not change during the whole growing process. These CFD modeling software (COMSOL Multiphysics) built-in library.

### Agent-Based Model (ABM) of cells growing on microcarriers

In order to study how the mechanical stress, determined by the stirred tank operative conditions, affects cell biomass, a three-dimensional agent-based model of cells growing on microcarriers moving in the fluid media of the stirred tank was developed. The model was implemented using the Biocellion agent-based modeling framework[11]. In the ABM formulation, each cell and microcarrier is represented as a spherical agent that is propelled by forces calculated from the computed velocity field of the fluid media. By including the cells and microcarriers explicitly, the model allows us to study the relationships between mechanical interactions and important biological processes such as cell proliferation and cell death.

#### Equations of motion

Newton’s equations were used to model the dynamics of both cells and microcarriers [13]:

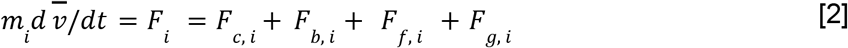

Where

- *m* is the mass of an agent *i* representing a microcarrier or a cell
- 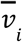 is velocity of the agent *i*
- *F*_*i*_ is the total force exerted on agent *i*
- *F*_*c,i*_ is the total force exerted on agent *i*due to contact with other agents
- *F* _*b,i*_ is the force on agent *i*due to contact with the bioreactor boundary
- *F* _*f,i*_ is the total force exerted on agent *i*due to fluid flow, this is the so-called drag force
- *F* _*g,i*_ is the force due to gravity and buoyancy

The mass *m*_*i*_ is constant for microcarriers and changes during the proliferation cycle for cells as described in equation [9]. The mass values and ranges can be found in S2 Table. The contact force on an agent *i* is the sum of all mechanical forces that act on agent i due to interactions with other agents and with the bioreactor boundary:

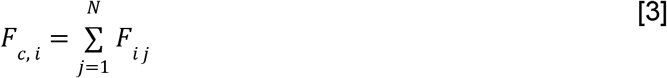

where *F* _*i*,j_ is the force exerted by other agents (cells and microcarriers) that form a bond with agent *i*. A bond is created between two agents when the distance between their centers becomes smaller than a threshold value δ_*c*_. Similarly, a bond between two agents is broken when the distance between their centers becomes larger than δ. The force between bonded agents is treated as a spring-bound system, and is described by the following equations:

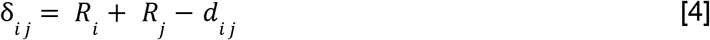

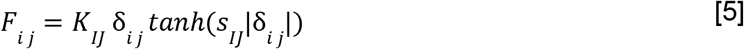

where *d* _*ij*_ is the distance between the centers of agents *i*and *j. I* and *J* represent the type (cell or microcarrier) of agent *i*and *j*, respectively. The bond between two agents, with the parameter *KK*_*IJ*_ being the spring constant of the bonds, is an attractive force when the distance is greater than (*R* _*i*_+ *R* _*j*_) and a repulsive force when the distance is less than (*R* _*i*_+ *R* _*j*_). The attractive force between a pair of agents grows with distance until the bond breaks for distances greater than *f*_*IJ*_ (*R*_*i*_+ *R* _*j*_) after which the agents become unassociated. The stiffness of the bond between two agents is controlled by the parameter *S* _*IJ*_ See [14] for a more detailed description of the bond model.

As an approximation for cell deformation when adhering to the microcarrier while using rigid spherical agents, the cell division radius *R*_*div*_ was subtracted from the microcarrier radius as given in S1 Table, resulting in the simulated microcarrier radius *R*_*m*_ as given in S2 Table. This correction places the cell centers approximately at the original microcarrier surface, making the amount of cells that fit on the microcarrier surface more realistic.

The forces on agents due to contact with the bioreactor boundary is modelled with a repulsive interaction force that is proportional to the overlap δ _*b, i*_ between the spherical agent *i*and the bioreactor boundary:

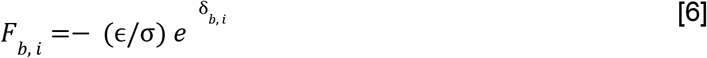

where ϵ captures the magnitude of the interactions between agents and boundary, and σis a scale factor of the order of the agents’ sizes [15].

In this work, the fluid drag force is based on Stokes flow past a sphere, and it is given by:

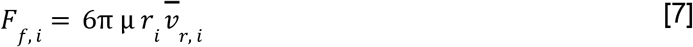

where μ is the dynamic viscosity of the fluid, *r*_*i*_ the radius of agent i, and 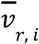 is the relative velocity of the agent i with respect to fluid velocity [16]. The fluid velocity of the flow is interpolated from the CFD velocity field as a weighted average of the velocities at the four vertices in the CFD grid that form the smallest tetrahedron containing the center of the agent. The relative weight for each vertex is the volume of the sub-tetrahedron formed by replacing just that vertex with the agent’s center.

Finally, the effects of gravity and buoyancy on agents are represented by the following equation:

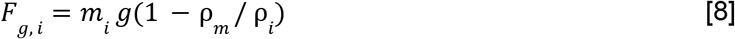

where *g* is the standard acceleration of gravity, ρ_*m*_ is the density of the medium, and ρ_*i*_ is the density of the agent.

The position of the agents was calculated from the force every time step using a Verlet leapfrog integration algorithm.

#### Cellular Phenotypes

Previous studies suggested that mechanical stress induces cell death, thus limiting the number of cells supported by a microcarrier [12]. However, there are other possible factors that limit cell growth such as cells entering a stress induced quiescent state instead of cell death. To study the different causes of reduced cell growth on microcarriers we consider cell proliferation, cell quiescence, and cell death (Fig. 4). Moreover, we model cell state transition due to external mechanical stress only, determined by mechanical stress threshold, σ_*D*_ and σ_*P*_ (Fig. 4). The following subsections describe in more detail the three cellular processes included in the model.

**Fig 4.**
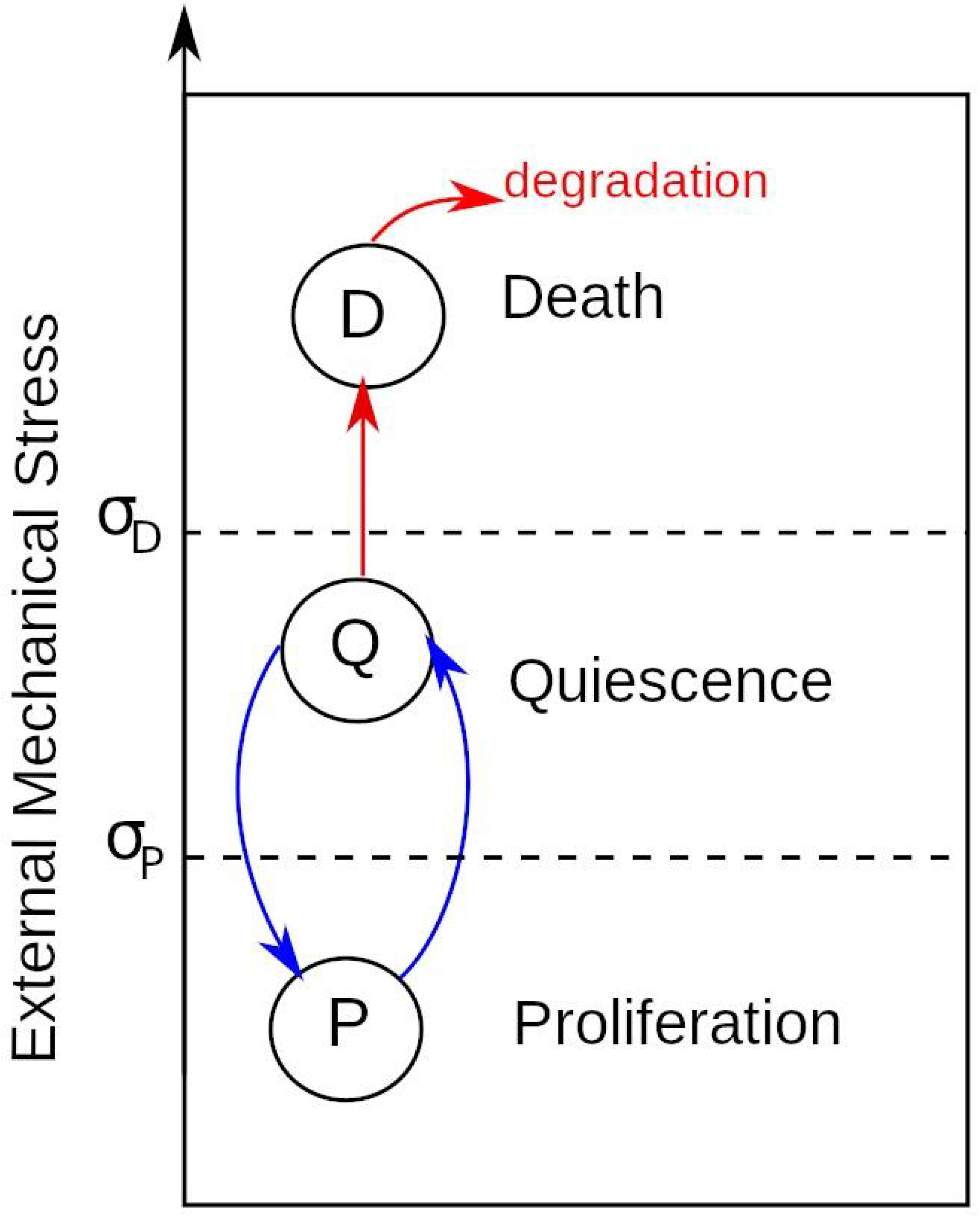
Phenotypes and their transitions due to external mechanical stress.

Under our current assumption of abundant nutrients, Hill-type stress based modulation of the growth rate of cell *i* was used to model cell growth, :

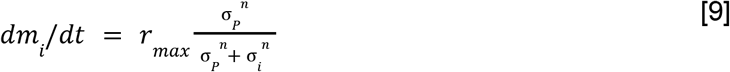

where *r*_*max*_ is the maximum proliferation rate obtained directly from the doubling time. This doubling time is for the sake of coupling to the fluid dynamics model, artificially scaled to ten doublings per second. This enables a direct strong coupling of the dynamical movement of microcarriers and cell experience of fluid forces to the proliferation of the cells. See the discussion section for an explanation of this approximation. σ_*i*_ is the mechanical stress on cell *i*, σ_*P*_ is a parameter that modulates the effect of mechanical stress on growth rate, and *n* is an integer that modulates the steepness of the Hill-type equation. Equation [9] dictates the transition between the normal proliferative cell state and the quiescent state where no proliferation occurs. Cells will stay in quiescence until the experienced stress decreases again below the threshold.

To compute the mechanical stress we first need to compute the tensor stress (*S*_*i*_) of agent *i*, due to interactions with other agents [17]:

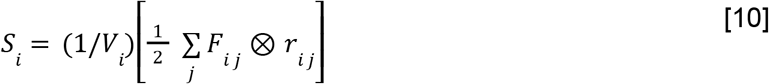

where ⨂is the tensor product of the two vectors, *F*_*ij*_ is the force and *r* is the distance vector between agent centers. The volume of the cell (*V*_*i*_) is computed assuming the cells are of constant density ρ, *V*_*i*_= *m*_*i*_/ρ. From the volume, we compute the new radii of cells. The mechanical stress used to modulate cell growth is the average of the principal stresses computed as the trace of the stress tensor:

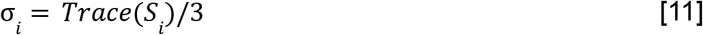

A cell division event is performed when the cell radius is greater than a user defined threshold *R*_*div*_. In cell division, a cell *i* is replaced by two daughter cells, one daughter has a mass between (*m*/2 − 0. 1*m*) and (*m*/2 + 0. 1*m*) (a random value is drawn from a uniform distribution between these two limits) and the other takes the remaining mass to ensure that the total mass is conserved. The two daughter cells are placed in the plane tangential to the microcarrier (sphere) that passes through the center of the mother cell. Only cells that are attached to a microcarrier divide. The direction within the plane in which the two daughter cells are placed is randomly selected. Both daughter cells are placed a distance (*R*_1_+ *R*_2_) /4 from the center of the mother cell, where *R*_1_ and *R*_2_ are the radii of the two daughter cells.

The model of cell death was adapted from J.A. Bull et al. [18]. Cells that are subjected to high mechanical stress die if they remain under high mechanical stress for longer than a threshold time *T*_*D*_. To determine the non-viable cell population induced by high mechanical stress we test if σ_*i*_> σ_*D*_, where σ_*D*_ is a mechanical stress threshold. Each cell is assigned a stress time counter τ_*i*_, which evaluates when a cell is under mechanical stress and evolves as follows:

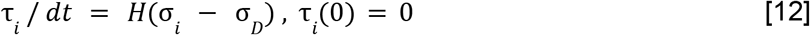

Where *H*(*x*) is the Heaviside step function (1 if x >= 0 and 0 otherwise). Cells transition to a cell death state when τ_*i*_> *T*_*D*_. After a cell transitions to a cell death state it enters into a disintegration process in which the cell biomass decays according to:

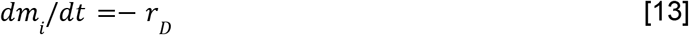

The cell is removed from the overall population when the cell radius is smaller than 10 micrometers. Moreover, *r*_*D*_ was setup to 50% of maximum proliferation cell death (*r*_*max*_).

### Sensitivity analysis

To investigate the model sensitivity with respect to its input parameters we performed a sensitivity analysis according to the method of Morris [19] with additional improvements by Campolongo et al. [20]. This method is also known as the Elementary Effects Method and is widely used to get an impression of the sensitivity of a model that has many parameters and long simulation times [21].

This method is well suited when the high number of input parameters makes the more thorough variance based techniques infeasible. It can be used to rank parameters according to their influence on the model output. The method as implemented in the SALib library was used to perform the analysis [22].

The sensitivity analysis was performed for the indicated biomechanical parameters in S6 Table since these are considered as the relevant ABM modeling parameters. Sensitivity analysis of the CFD parameters and the physical system was not done in this project, but might be explored in future work.

Sampling the parameter space using Morris’ sampling method was done for the indicated parameters from the ranges as indicated in S6 table using the Elementary Effects sampling method. These ranges were inspired by conversations with the industry and literature and were specified to cover a physically realistic extent. The sampling was conducted with r = 10 trajectories and p = 4 sampling levels. Some parameters were sampled from an exponential distribution, this is also indicated in S6 Table.

## Results

### Fluid characteristics for different impeller rotation speeds (CFD)

The steady-state flow behaviors (velocity magnitude, fluid shear stress, and Kolmogorov length) at impeller rotation speeds of 60 and 220 rpm are illustrated in Fig 5. These modest and vigorous rotational speeds induce similar fluid flow patterns but quantitatively different flow metrics. The Reynolds number has been calculated as 1430 for 60 rpm and 5420 for 220 rpm considering

**Fig 5.**
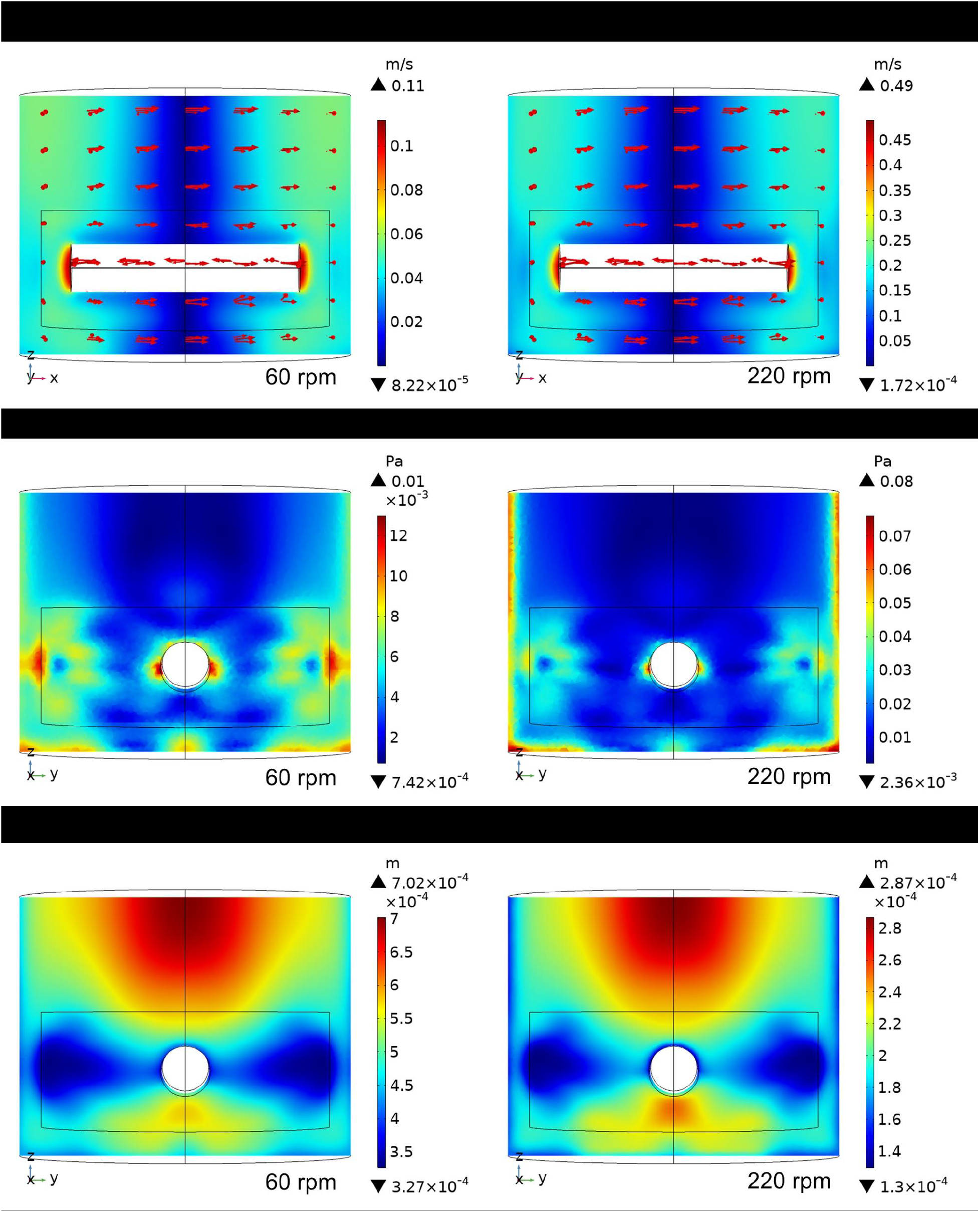
Computational fluid dynamic simulations at 60 rpm and 220 rpm. Fluid velocity profile (top), fluid shear stress (middle) and Kolmogorov eddy length scale (bottom).

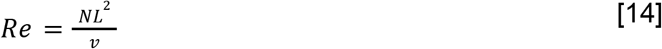

where *N* is the impeller rotations per second, *L* is the length of the impeller, and *v* is the kinematic viscosity [23]. One can see from the vertical slice cut parallel to the x-z plane that at both rotor speeds the velocity magnitude is largest near the tips of the impeller and lowest in a vertical cylindrical region at the center. However, as is evident by the relative ranges on the color-coded legends, the velocity magnitude at 220 rpm is about four to five times that at 60 rpm.

The shear stress is a parameter, also displayed in Fig 5, that influences the biological processes in bioreactors. Similar to the velocity magnitude profiles, the shear stress shows comparable patterns at 60 and 220 rpm. Generally, higher shear stress exerts more influence on the cells, and is observed around the tips of the impellers, similar to the results reported in Ghasemian et al.[24]. That work simulated the hydrodynamic characteristics within a spinner flask at rotational speeds of 40, 60, 80, and 100 rpm, where the shear stress ranged from 0 to about 80 mPa and were similar to the shear stress produced in this study at 60 rpm. Heathman et al. [25] studied the effect of agitation rate on the growth of bone marrow-derived human mesenchymal stem cells (BM-hMSCs) on microcarriers in a stirred-bioreactor. The study found that, although microcarriers and cells were not entirely suspended at low speed (80 rpm), the highest number of post-harvest cells was achieved at 80 rpm. The number of cells decreased with agitation rate to about 20% for 225 rpm. In this study, a much higher rotational speed (220 rpm) was also simulated to show the pronounced negative effect of elevated stir speeds on cell growth.

Kolmogorov length is another parameter that helps understand the influence of turbulent eddies on cells [26]. The Kolmogorov length or scale theory states that small-scale motion is a function of the dissipation rate per unit mass and the kinematic viscosity [27]. The Kolmogorov scale is typically represented as:

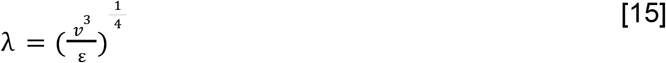

where λis the characteristic size of the eddies, *v* is the kinematic viscosity, and ε is the rate of dissipation of turbulent energy [28].

Although the two rotational speeds in this study show similar Kolmogorov length distribution patterns within bioreactors, their Kolmogorov length values are considerably different. Both rotational speeds have lower Kolmogorov length values around the tips of impellers. The Kolmogorov length values of the 60 rpm scenario in the whole bioreactor are larger than the microcarrier size of 180 μm; while that of the majority regions of the 220 rpm scenario are much smaller than or similar to the microcarrier size. These simulated Kolmogorov length values are also similar to that reported in Ghasemian et al.[24]. Croughan et al. [12] reported that the relative cell growth rate significantly decreased when the Kolmogorov length was smaller than 125 μm and detrimental effects appeared to come into play when the Kolmogorov length dropped below about 100 μm, which was about half of the average microcarrier diameter of 180 μm. In this study, the minimum Kolmogorov length is about 60 μm, which is much smaller than the critical length, indicating potential significant damage to the cells when microcarriers flowed into these regions.

### Multiple growth behaviors induced by mechanical stress (ABM)

Results from simulations of cell growth on a single microcarrier in the absence of fluid movement are presented in this section. These simulations will help to study how cell state transition and growth are affected by interactions between cells, interactions with other cells and those with the microcarrier. The parameters of the simulations are shown in S2 Table.

For each cell *i*, stress σ_*i*_ is computed as described in “Cell Growth subsection”. Compressive forces contribute positively and tensile forces contribute negatively to the overall mechanical stress sensed by cells. Fig 6A shows a simulation snapshot of cells growing on a microcarrier and Fig 6B shows a histogram of mechanical stress experienced by cells. The color of cell *i* represents the magnitude of mechanical stress. Most of the cells in this example experience compressive forces exerted by their neighbors as indicated by their reddish color (Fig 6A) and by the predominantly positive values for net stress on a cell reported in the histogram (Fig 6B). Literature reports note the prevalence of compressive forces in cell monolayers bound to flat substrates [29] and those within circular confinements [30,31]. This figure suggests the mechanical stress sensed by the cells on the microcarrier may be rather variable even in the absence of shear forces. It is notable, however, that the distribution of stress profiles experienced by the cell population on the microcarrier is semi-normal (Fig 6B).

**Fig 6.**
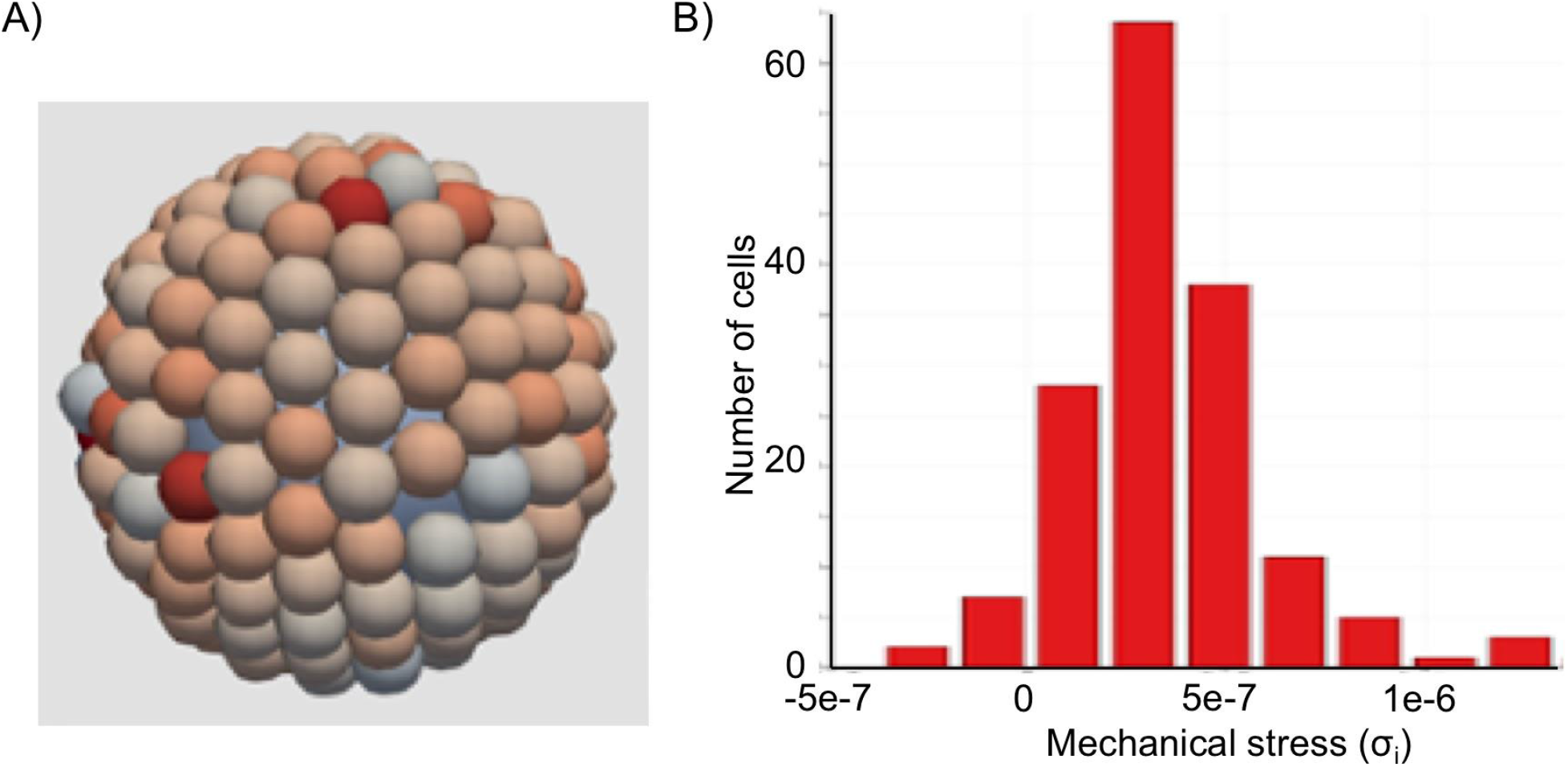
Agent based model of cells on a microcarrier. (A) Cells growing on top of a microcarrier. Red indicates compressive stress and blue indicates tensile stress. (B) Histogram of mechanical stress sensed by the cells growing on the microcarrier.

First, a scenario in which mechanical stress does not induce cell death but reduces the capacity of cell proliferation was tested, see equation [9]. Within our modeling approach, this scenario is simulated by setting σ_*D*_ = 1, a very high value according to Fig 6B. Fig 7A shows the number of cells growing on the microcarrier as function of time for different values of σ_*P*_ and *n* = 5, while similar results for other values of *n* are shown in the Supporting Material, S3 Fig. The results in Fig 7A show different growth trends for different values of σ_*P*_. For high values (σ_*P*_ = 1*e* − 6) the cell count first grows exponentially and then deviates from exponential growth; however it does not reach a constant value. For smaller values of σ_*P*_, the number of cells reaches a constant value. A second scenario was simulated, in which mechanical stress does not change proliferation rate but induces cell death, thus limiting the population size. This scenario is simulated by suppressing the transition of cells to the quiescent state, that is making σ_*D*_ = σ_*P*_. Fig 7B shows the cell count as a function of time for different values of σ_*D*_ and *T*_*D*_ = 0. 1τ_*d*_, while results of other values of *T*_*D*_ are shown in S4 Fig of the supporting material. Similar to Fig 7A, Fig 7B shows different growth trends for different values of σ_*D*_ For σ_*D*_ = 1*e* − 5 μ*N*/μ*m*^2^, the population growth is exponential with the same rate as the unconstrained exponential growth (σ_*D*_ = 1 μ*N*/μ*m*^2^). For smaller values of σ_*D*_ the population reaches a constant value similar to Fig 6B. It is worth noting that initially, the number of cells grows at the same rate as in the unconstrained growth for the scenario in Fig 7B, whereas the cells can grow at much slower rates for the scenario of Fig 7A and S3 Fig.

**Fig 7.**
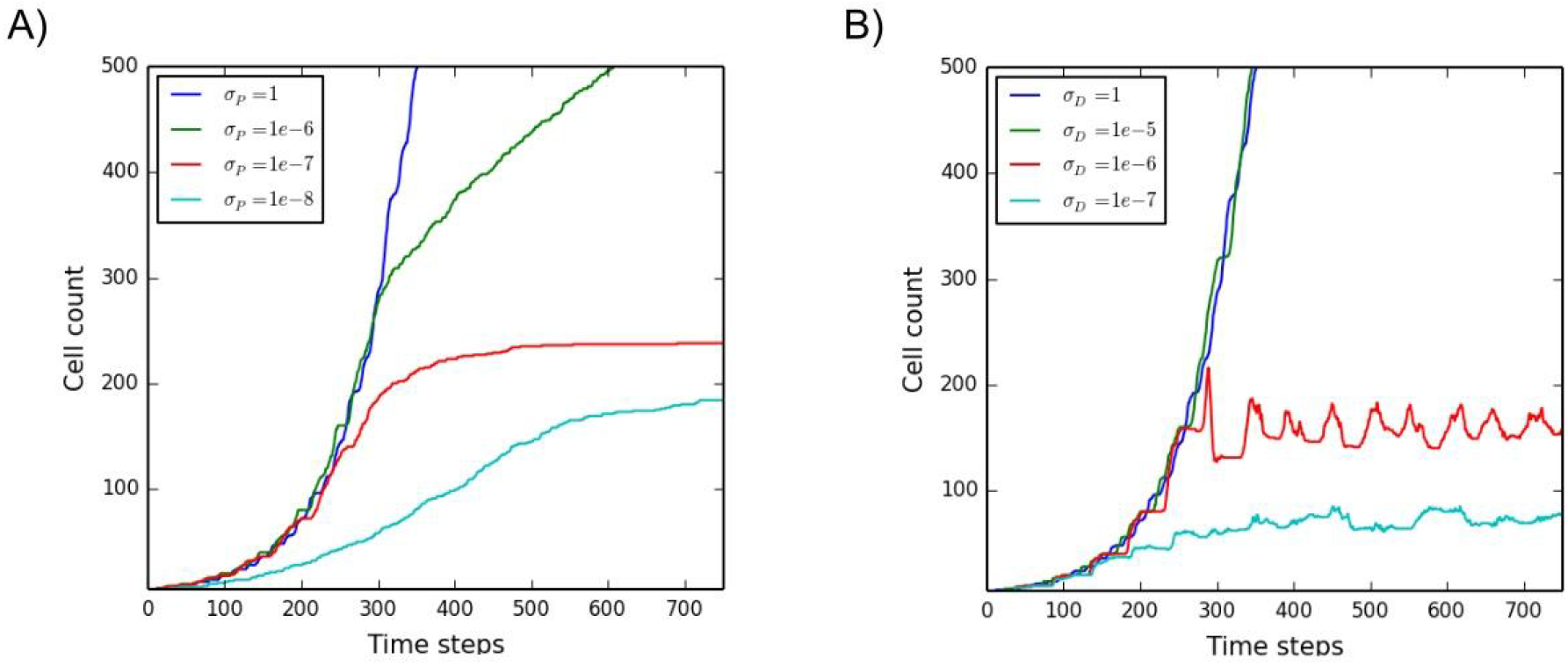
Cell population on a single microcarrier as a function of time. The amount of cells per microcarrier over time for (A) various values for the proliferation threshold and (B) various values for the cell death threshold

### Impact of impeller rotation speed on microcarrier distribution (CFD and Lagrangian Particle Tracking)

To assess the distribution of microcarriers throughout the bioreactor at various times with various rotation speeds, simulations were performed with 1000 microcarrier agents without cells growing on them. The velocity fields from the CFD simulations were used to perform Lagrangian Particle Tracking (LPT) using only the mechanical interactions of the Agent Based Model as described in equation [2]. Starting from a random position, the locations of all microcarrier agents were accumulated over 200000 simulation time steps to give a relative distribution of microcarriers as shown in Fig 8. It can be seen that at early simulation times (e.g. 120000 simulation steps), the initial random distribution is still prevalent. However from 500.000 simulation steps onward, high density zones become apparent, growing in intensity for longer simulation times as the randomness of the initial conditions disappears. A clear high density spot appears just above the impeller tip for impeller rotation of 60 rpm, as can be seen from Fig 5 this corresponds to a high stress region. Another high density zone can be seen in the top right corner for the 220 rpm simulations, this zone grows in intensity beyond the scale depicted in the figure colormap.

**Fig 8.**
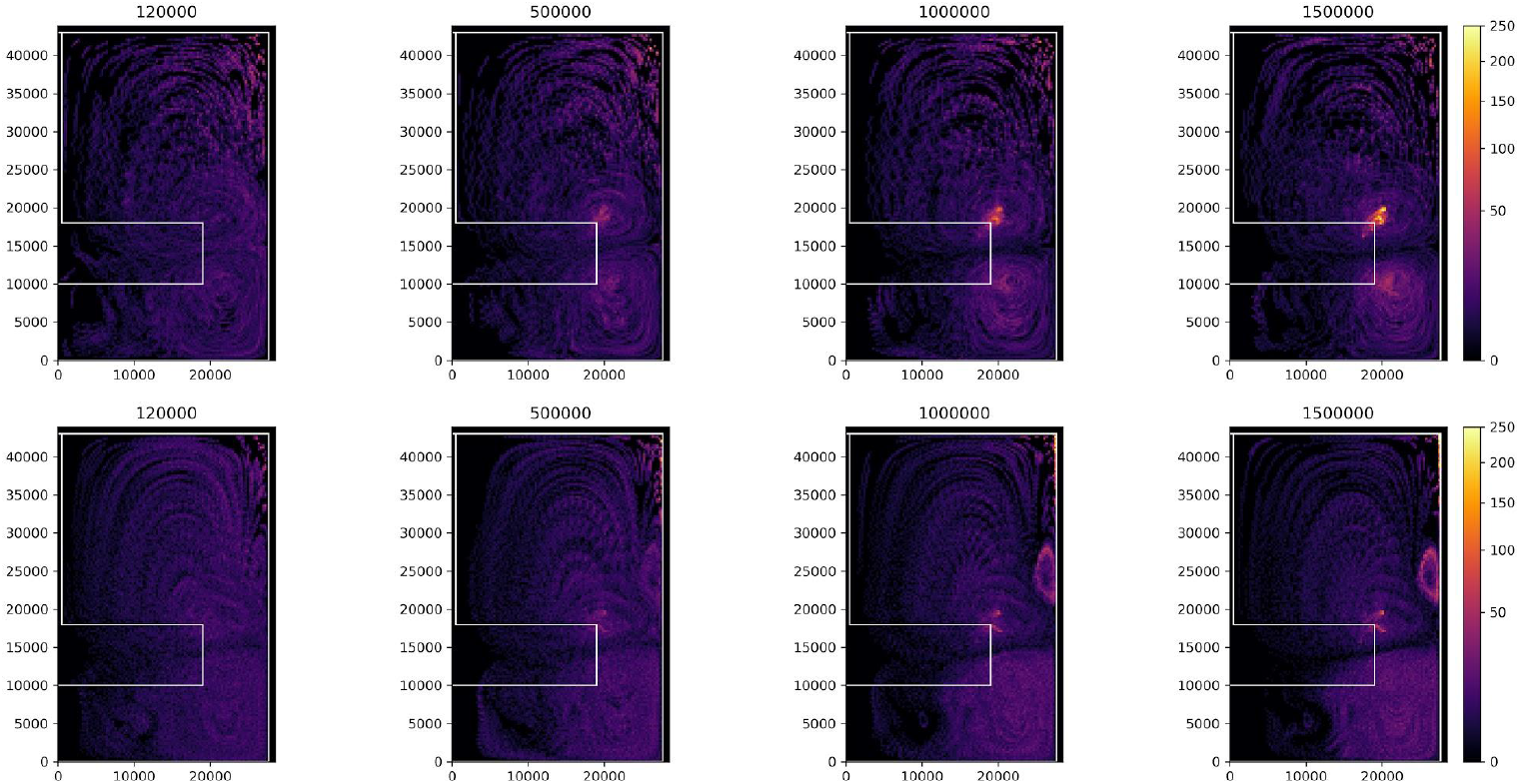
Microcarrier density heat maps due to mechanical forces. The count of microcarriers, accumulated over the whole bioreactors radial symmetry and 200000 simulation time steps in a bin size of 275 *μm*^2^. The top row shows simulations with the 60 rpm velocity field and the bottom row uses the 220 rpm velocity field with the high end of the simulation step accumulation range given at the top of each column.

### Impact of impeller rotation speed on cell growth (CFD and ABM integration)

The ABM and CFD methods were combined to simulate how the mechanical stresses inside the stirred tank influence the biomass accumulation of cells attached to spherical microcarriers. In our model, fluid flow influences how individual cells move, grow and proliferate. Because in CFD simulations the modeled bioreactor reaches its steady-state flow in just a few seconds, transient flows in the whole-system simulation were ignored. As described in Methods section, the distribution of fluid velocities at steady state to compute the drag forces *F*_*f, i*_[Eq. 7] was used. For these simulations we explored several values of parameters σ_*D*_ and σ_*P*_, while other parameters are specified in S2 Table. Moreover, we observed that the fluid can produce cell detachment specially at 220 rpm, thus we report only live cells that are attached to the microcarrier.

Biomass levels for simulations at both 60 rpm and 220 rpm were compared. These simulations test whether the model can qualitatively recapitulate the trends observed experimentally by Croughan *et al*.[12], described in Methods. They observed that the biomass level in the bioreactor decreases with increasing impeller rotation rate, and that high rotor speeds (220 rpm) induce cell death (Fig 2).

Fig 9 shows how the number of live cells (solid lines) change in time at 60 rpm and 220 rpm. Fig 9A and Fig 9B show simulations for σ_*D*_ = 10^−6^, σ_*D*_=10σ_*P*_ and σ_*D*_= 10^−7^, σ_*D*_= σ_*P*_, respectively. Other values of σand σwere also tested; the results from these simulations can be found in the Supporting material, S5 Fig. As expected, in all the simulations the number of cells at 220 rpm grows slower than that at 60 rpm. This is observed even when the number of dead cells is zero (Left hand side of Fig 9A). Moreover when σ_*D*_ =σ_*P*_, the cells at 220 rpm grow very slowly but due to regions of high mechanical stress the number of cells went to zero. These simulation results are in good qualitative agreement with those reported by Croughan *et al*.[12]; increasing rotor speed reduces biomass growth rates and cell death is induced at very high rotational speeds.

**Fig 9.**
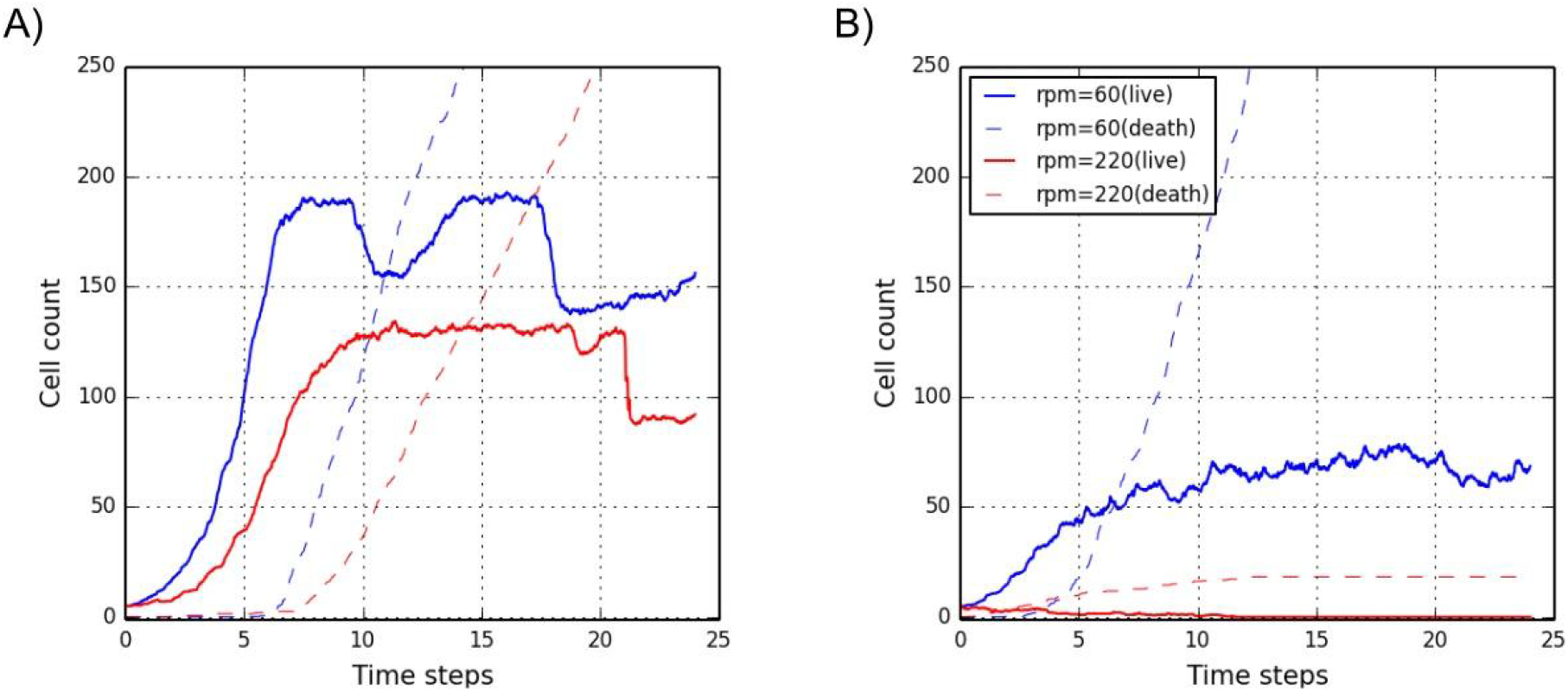
Cell population on a single microcarrier as a function of time for two rotor speeds, 60 and 220 rpm. With cell death thresholds of σ_*D*_ = 10^−7^ and (A) σ_*D*_ = 10σ_*P*_ and (B) σ_*D*_ = 100σ_*P*_. Solid lines represent live cells attached to the microcarrier and dashed lines represent the cumulative number of death cells. The cell counts are averages over 5 simulations; and one time step in this case represents the theoretical doubling time of cells.

### Sensitivity analysis results

For every parameter sample, ten replicates of the simulation were run and the average output taken for analysis. The output of the model that was considered in the sensitivity analysis was set to the value of the number of cells attached to a microcarrier at the 20000th and at the 50000th time step. The latter time step was chosen since many configurations as shown in S5 Fig yielded a steady state at or around that time step. This choice of output potentially influences the measured sensitivity of the model. However no significant changes in the parameter ranking were found by using the output at later time steps. The results of the sensitivity analysis change significantly when choosing an earlier timestep.

The results of the sensitivity analysis are shown in Fig 10 and indicate that after 50000 timesteps the highest variation in the output is due to a difference in the parameter that determines the cell-cell bond strength (A_AGENT_BOND_S_CC). However the uncertainty σin the elementary effect is large compared to the elementary effect itself.

**Fig 10.**
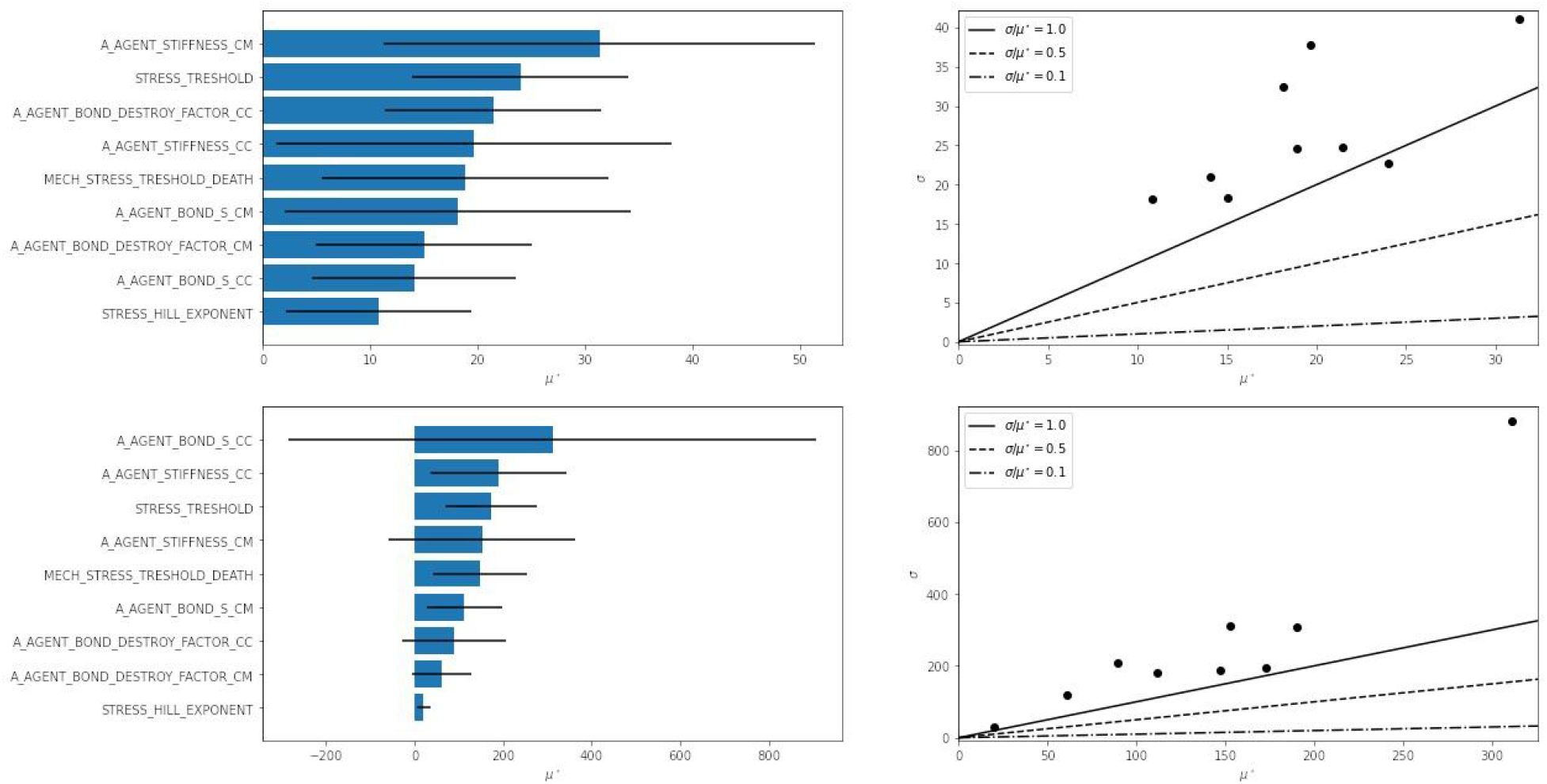
Results of sensitivity analysis. Parameters ranked by their elementary effect μ* on the model outcome (left) and the elementary effect plotted against the deviation σ of that elementary effect (right). The sensitivity was analyzed for the number of attached cells at simulation step 20000 (top) and 50000 (bottom) steps. See S6 Table for the description of parameter names.

After 20000 steps the results show a stronger dependence on the cell-microcarrier bond stiffness (A_AGENT_STIFFNESS_CM) and the stress threshold (STRESS_TRESHOLD). The uncertainty in elementary effects is also considerably smaller.

## Discussion

Combining multiple modeling approaches in a more comprehensive model can generate novel insights and identify emergent properties in the system under study. Indeed, CFD and ABM have been used together in several diverse multiscale models that simulate disaster responses [32], cell and particle migration through blood vessels [33–36], and movement of zooplankton in complex flow environments [37]. Notably, CFD and ABM *per se* have never been used together to understand the process dynamics within a bioreactor relevant to cultivated meat. However, combined models for biological and physicochemical dynamics have been used to examine different parallel-plate bioreactor configurations for growth of tissue [38]. Here, a unilineage model was employed to describe the replication and differentiation of stem cells, and the physicochemical processes were modeled by the Navier-Stokes and convective-diffusion equations.

Despite the advancements made in prior studies, it is worth reiterating that the present work aims to prototype a new computer modeling methodology for adhesive cell cultures in bioreactors. The new framework was used to qualitatively recapitulate the empirical and theoretical results from Fig 2 [12]. Several assumptions and simplifications were made in the novel model:

I. Assumption. The only impact of mechanical stress on cell biology is to slow cell proliferation rate and induction of cell death.
II. Assumption. Capturing the relevant fluid dynamics behaviors is accomplished at a much shorter simulation time-step than is needed to capture the biological response behaviors of cells.
III. Simplification. Biomass accumulation and microcarrier count does not significantly affect the fluid properties.
IV. Simplification. Cell growth is independent of the concentration of molecules (such as nutrients, oxygen, or H^+^ ions) in the media.
V. Simplification. Microcarrier and cell densities are smaller than those used and observed in laboratory experiments.
VI. Simplification. By using a RANS approach for CFD we do not model small turbulent eddies explicitly, therefore these eddies are not taken into account when calculating the mechanical stress.
VII. Simplification. The effect of the fluid on microcarriers and cells was modeled using the Stokes law for drag force.
VIII. Simplification. The Stokes drag force was calculated for every agent (cells and microcarriers), and in this calculation no shielding by other agents was taken into account.
IX. Assumption. The time dependent effect of the fluid on the cells will average out in steady state flow, for multiple microcarriers and multiple simulations.

Previous theoretical studies, specifically Croughan et al. [12], assert that cell death is the main driver of biomass reduction at higher rotor speeds but without providing experimental validation. Our understanding from experimentalists and the literature is that dead cells are difficult to count accurately because they may disintegrate into the surrounding media during the experimental period evading measurement post-experiment [39]. Assumption I introduces the alternative mechanism that mechanical stresses suppresses cells’ proliferation rates. Results in this study (Fig 8) support the hypothesis that this mechanism is also responsible for the overall reduced biomass level observed at higher rotor speeds. The ABM approach, facilitating representation of individual cells and the induced mechanical stresses they experience, thus provides a direct means to answer the question: “Instead of cell death, could a reduction in proliferation rate caused by mechanical stresses due to fluid flow on some cells explain the reduced biomass at higher stir speeds?” Answered in the affirmative for virtual experiments, there is now a stronger case for asking the same question of laboratory experimentalists.

It is notable that the use of the virial stress formulation (Eq. 11) for spherical cells can give rise to large fluctuations in this measure [40]. However, this work has continuously accounted for volume change in determining the tensor stress on each agent (Eq. 10), which is more accurate than a constant volume assumption in the application of virial stress determinations for atomistic stress [17]. Despite the apparent variability (Fig. 6A), the mechanical stress distribution for cells on the microcarrier is semi-normal (Fig. 6B), which mirrors the influence of compressive stress in tumor spheroid environments [40]. Furthermore, cells growing on the surface of microcarriers typically grow in a single layer which results in lower virial stress fluctuations per cell overall than in a spheroid system.

Assumption II and Simplifications III and IV simplify integration of ABM and CFD in this early modeling approach. Simplification III implies fluid properties are independent of the cell and microcarrier population. A CFD simulation performed on a bioreactor with only media therefore suffices to predict fluid velocities. Simplification V means both that cell ingestion and secretion rates as well as mixing are irrelevant, so that simulation can be performed on a homogeneous fluid. Because steady state is reached in under ten seconds of bioreactor operation, Assumption II permits a steady-state fluid flow to be used at all biological time-steps.

One advantage of this approach is that fluid dynamics can be modeled independently of the cell behavior as a stage whose output becomes an input to the ABM simulation. There is not yet a need for a feedback edge that would require recomputing CFD. However, all four of these simplifying assumptions are insufficient to bridge the discrepancy in biomechanics and biological time-scales. Namely, that a cell can move from one end of the bioreactor to the other in seconds, whereas its division into two cells takes about a day. Furthermore, simulating the millions of microcarriers and billions of cells in even small bioreactors, while possible using supercomputers, is beyond the computational capability of the desktop computers and small cloud clusters we have readily available. The workarounds are Assumption IX, to perform simulation using unrealistic proliferation rates in the time scale of seconds, and Simplification V, to use a number of microcarriers considerably smaller than typically used in experiments. The qualitative trends observed in Figs 7 and 8 are not expected to change due to these simplifications. Further model refinements will be needed to model the whole bioreactor system at realistic temporal and spatial time scales.

Assumption VI, although discrepant with the to be tested eddy length hypothesis mentioned by Croughan et al [12], allowed for simulating the whole bioreactor domain. Doing this with Direct Numerical Simulation or Large Eddy Simulation with a mesh on the scale of the individual cells would have made the simulation unachievable due to the large domain. Current efforts of the research group focus on coupling CFD models on different spatial scales and coupling ABM and Lagrangian particle tracking methods on different scales to overcome the (temporal and spatial) scale problem that led to assumption II and IV and to be able to simulate the direct effect of small scale turbulent eddies on the cell biology. This coupling approach would separate the spatial scales, where the position-dependent large scale flow profile around a microcarrier is tracked with Lagrangian particle tracking or a simple ABM model as it circulates through various velocity fields simulated on a large scale in a stirred tank. These large spatial scale flow profiles can then be fed into small spatial scale simulations that model the effect of small scale turbulence (CFD) on cells (ABM) explicitly. These small scale simulations can be used as statistical representatives to inform the large scale model with respect to several fluid and particle characteristics. A separation of temporal scales in the small spatial scale simulation can be achieved by running the CFD simulations for short intervals with small (sub-second scale) time steps at every (hour scale) ABM step.

Simplification VII was chosen as an approximation for calculating the drag force on both cells and microcarriers. This approach was taken as a similar approach has been used and validated in other microcarrier modeling efforts as well [41]. This simplification also implies that hydrodynamical lift forces (e.g. Magnus effect) were not taken into account. We take this to be a reasonable assumption since the agents are spherical bodies that move freely with the fluid with small velocity relative to the fluid and no significant spin.

Simplification VIII allows the use of a single force to calculate both microcarrier movement and agent stress. Rather than calculating the total drag force of a packed microcarrier, which would have been more accurate for the microcarrier movement, we chose this approach to be able to simulate the effect of fluid velocity gradients on the cells attached to microcarriers.

With regards to the sensitivity analysis, overall it seems that the uncertainties σ of the elementary effects are large. This can probably be solved by running more or larger (more microcarriers) simulations per parameter set so the output will statistically be more steady. However as a first exploration of parameter space the sensitivity analysis gives essential information for further research. It is notable that the strength and uncertainty of the elementary effect of the cell-cell bond strength parameter becomes very high while the other effects remain ranked in a similar order with comparable variance. A variance based exploration of the top three or four parameters with more simulations per parameter set will therefore be a logical next step.

In spite of the assumptions and simplifications, this unified CFD and ABM model was able to recapitulate the experimental observation that increasing rotational speeds in stirred-tanks reduces biomass growth rates. Thus, this model promises to offer an enhanced understanding of microcarrier-based stirred tank bioreactors that was not possible with the analysis conducted by Croughan *et al*. [12]. The damaging effects due to formation of turbulence are of special interest for further investigation. Researchers investigating turbulence have noticed eddy size determines its effect on solid particles [42–44]. Their observations suggest that microcarriers are harmlessly swept away by eddies whose curvature is smaller than the radius of the microcarrier, but that eddies adjacent to a microcarrier of sharper curvature instead exert distortional forces. The eddy-length model specifically proposes that damage occurs to cells when the Kolmogorov eddy length is below a critical value [12]. Given the single cell size spatial resolution that is possible with ABM and the turbulence modeling capability of CFD, this novel computer modeling methodology should make it possible to assess the validity of the eddy-length model theory. Croughan *et al*. [12] also noted that fluctuating fluid velocity components around a microcarrier change rapidly with time and position as it moves throughout a stirred bioreactor. These authors propose a “time-average” analysis model, where the position-dependent time-average flow profile around a microcarrier is tracked as it circulates through various time-averaged velocity fields in a stirred tank. Realistically, this is an oversimplification of dynamics within the bioreactor environment and neglects to capture how this fluid flow variability will manifest across a cell population. Here, the combined CFD-ABM model could offer more detailed information. Whereas previous modeling efforts for microcarrier cultures in stirred tank bioreactors used Euler-Euler or Euler-Lagrange approaches [41,45], and achieved results that are in agreement with experiments, these approaches are limited by the simplicity and/or continuous representation of the microcarriers and cells. When cell-cell interactions become more prevalent due to increased density, these approaches will not suffice. Using the combination of CFD with ABM enables cell resolution results and explicit addition of mechanical and biological interaction between cells and their environment. In this way the CFD-ABM approach can reveal detailed information regarding cell population heterogeneity, where this “cell-state spread” should be a proxy for bioreactor/bioprocess efficacy.

## Conclusions and future work

The combined modeling approach described in this paper proves to be effective for investigating the effect of various mechanisms in the bioprocess on several relevant scales, from full bioreactor fluid flow profiles and microcarrier distributions to single cell experience of mechanical forces. Future research will focus on refining the coupling between CFD and ABM and the different spatial and temporal scales to make the overall framework better suited for simulation of larger bioreactors. Next steps in the whole systems model development include adding additional physical and biological mechanisms. For CFD these include aeration, heat and mass transfer and for ABM these include nutrient consumption, oxygen utilization, cell deformation, gene regulation and cell signalling.

Beyond the addition of other physical and biological mechanisms, further work must also refine existing models such as the adhesion model for agents. In this case, the current tanh based adhesion model as described in the Methods has proven useful [14], but it is a non-realistic representation of the actual biomechanics. Ongoing investigations around this topic take into account several adhesion modeling approaches (JKR, Lennard-Jones, integrin based model) [40,46,47].

Another aim of ongoing work is to verify the implemented mechanisms and parameters of the modeling framework in a simple experimental configuration that can be controlled precisely. Such a configuration, the IBIDI computerized pump system (ibidi GmbH, Martinsried/Munich, Germany), has been identified and is being used in several labs by collaborators to refine the effect of various shear stress time profiles on proliferation rate, migration, detachment and induced cell death. A further intent of this work is to translate as many relevant model parameters as possible to measurable experimental parameters so that experimental setups can be directly or nearly translated to the computer model. Additional verification studies should also refine the cell agent response to compressive and tensile profiles potentially using experimental results from monolayer stress microscopy [31].

Experiments to validate the whole system stirred tank model are also being set up in collaboration with cultivated meat companies and research groups that use these bioreactors. The ultimate intention is that each partner can examine specific parameters of the whole systems model using controlled experiments, after which the framework can be used to analyze and optimize their production set up. In this way the modeling framework is intended to replace many expensive lab experiments that these companies would have to run in order to test and optimize their large scale meat production facilities.

## Acknowledgments

This work is an initiative by the Cultivated Meat Modeling Consortium (www.thecmmc.org). The authors would like to thank the other members for engaging and informative discussions.

## Supporting information

**S1 Table.** Parameters obtained from Croughan et al., 1987 and other sources.

| Parameter | Value | Magnitude (units) | Source |
| --- | --- | --- | --- |
| Cell name | FS-4 |  | [12] |
| Cell origin | Human diploid fibroblast |  | [12] |
| Cell shape | - |  |  |
| Cell diameter | 18-22 | $\mu\text{m}$ | [48] |
| Cell volume | 3000-6000 | $\mu\text{m}^3$ | [48] |
| Cell mass | 3.50e-09 | g/cell | [6] |
| Microcarrier type | Cytodex-1 |  |  |
| Microcarrier diameter | 180 | $\mu\text{m}$ | [49] |
| # microcarriers/mass | 6.80e+06 | microcarriers/g dry weight | [49] |
| Swelling index | 18 | mL/g dry weight | [49] |
| Use density | 3 | g/L | [49] |
| Total # of microcarriers (in 100 mL) | 2.04e+06 | microcarriers | * |
| Microcarrier dry mass | 1.47e-07 | g/dry carrier | * |
| Microcarrier wet mass | 2.79e-06 | g/wet carrier | * |
| Growth temperature (GT) | 37 | degrees C | [12] |
| pH | 7.3 | - | [12] |
| Vessel geometry | 5.5 | cm (internal diameter) | [12] |
| Media volume | 100 | mL | [12] |
| Specific density (@ GT) | 1.01 | - | [12] |
| Density (@ GT) | 1.003233 | g/cm<sup>3</sup> | * |
| Media viscosity (@ GT)<br>(Newtonian) | 1 | cP | [12] |
| Media height | 4.21 | cm | * |
| 1/3 media height | 1.4 | cm | * |
| Stir bar length | 3.8 | cm | [12] |
| Stir bar diameter | 0.8 | cm | [12] |
| Agitation | 60,140,180,220 | rpm | [12] |
| Medium change (75 %) | 4th | day | [12] |
| Starting glucose concentration | 4.5 | g/L | [12] |

\*calculated

**S2 Table.** Parameters that were used in the models. The values in this table were used in all simulations where the parameter is not otherwise specified. The * symbol in the source column indicates that the parameter was manually calibrated to avoid numerical instability while keeping the agents within the bioreactor boundary. † indicates that this parameter was inspired by conversations with industry partners and experimental groups. (SA) indicates that this parameter was taken into account in the sensitivity analysis. Cell radii in this research were inspired by [48] but adjusted upward by 5 microns to account for a larger radius through cell spreading.

| Name | Symbol | Value | Source |
| --- | --- | --- | --- |
| Cell properties (ABM) |  |  |  |
| Density of cells | $\rho_i$ | 8.36e-16 kg/$\mu\text{m}^3$ | [6,48] |
| Cell division radius | $R_{div}$ | 15 $\mu\text{m}$ | [6,48] |
| Mechanical stress modulator | $K_s$ | $1\text{e-}7 \mu\text{N}/\mu\text{m}^2$ | (SA) |
| Microcarriers properties (ABM) |  |  |  |
| Microcarrier radius | $R_m$ | $75 \mu\text{m}$ | [6,48,49] |
| Microcarrier density | $\rho_m$ | $8.32\text{e-}17 \text{ kg}/\mu\text{m}^3$ | [49] |
| Fluid (water) properties (CFD) |  |  |  |
| Density | $\rho$ | $1.0\text{e-}18 \text{ kg}/\mu\text{m}^3$ | - |
| Viscosity | $\mu$ | $1.0\text{e-}9 \mu\text{N}\cdot\text{s}/\mu\text{m}^2$ | - |
| Bioreactor geometry (CFD and ABM, only the coupled model) |  |  |  |
| Bioreactor height | $H_B$ | $42895 \mu\text{m}$ | [12] |
| Bioreactor diameter | $D_B$ | $55000 \mu\text{m}$ | [12] |
| Impeller length | $L_I$ | $38000 \mu\text{m}$ | [12] |
| Impeller diameter | $D_I$ | $8000 \mu\text{m}$ | [12] |
| Cell-cell mechanical interactions (ABM) |  |  |  |
| Cell-cell spring constant | $K_{cc}$ | $1\text{e-}3 \mu\text{N}/\mu\text{m}$ | (SA) |
| Cell-cell bond flexibility | $s_{cc}$ | 0.2 | (SA) |
| Thresholding factor for cell-cell bond breaking | $f_{cc}$ | 1.1 | (SA) |
| Cell attachment to microcarriers (ABM) |  |  |  |
| Cell-microcarrier spring constant | $K_{cm}$ | $2.2\text{e-}3 \mu\text{N}/\mu\text{m}$ | (SA) |
| Cell-microcarrier bond flexibility | $s_{cm}$ | 0.2 | (SA) |
| Thresholding factor for cell-microcarrier bond breaking | $f_{cm}$ | 1.1 | (SA) |
| Cell bioreactor interactions (ABM) |  |  |  |
| Magnitude of boundary repulsive forces | $\varepsilon$ | $2.0\text{e-}9 \mu\text{N}$ | * |

| Initial conditions for simulations (ABM) |  |  |  |
| --- | --- | --- | --- |
| Initial number of cells per microcarrier |  | 5 | † |
| Initial cell radius | | 11 - 14.5 $\mu\text{m}$ | [48] |

**S3 Fig.**
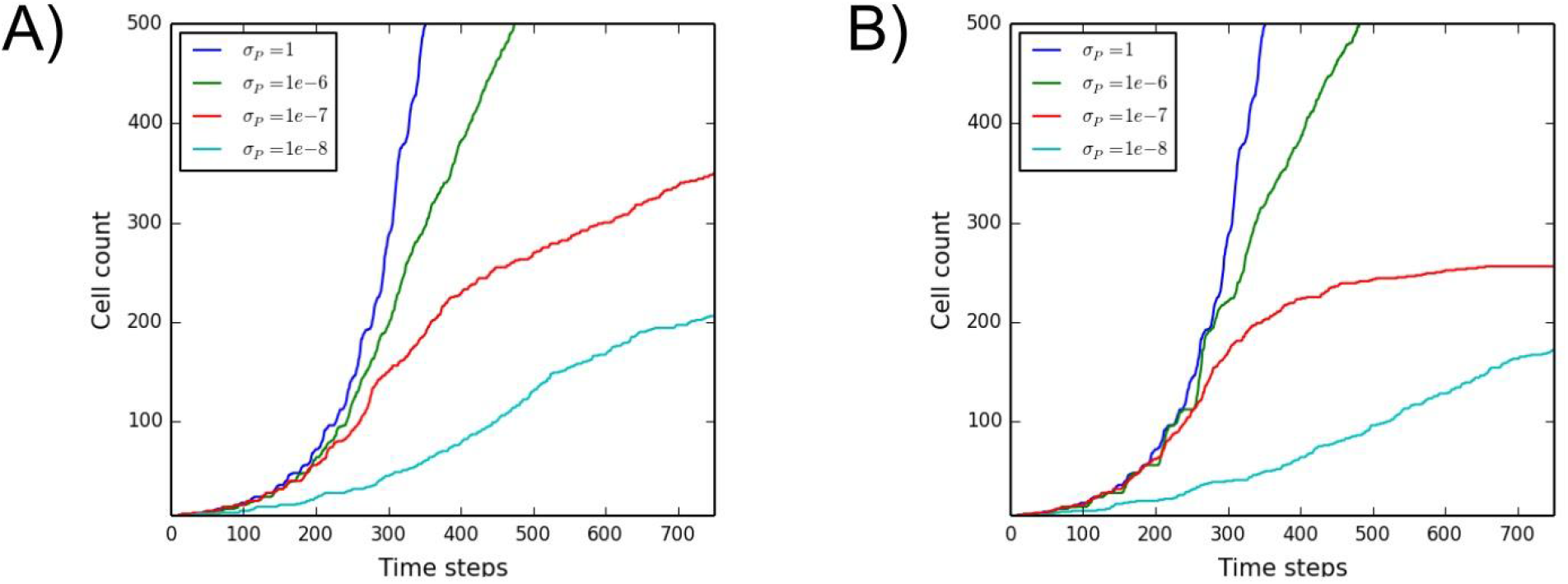
Cell population as a function of time without CFD effects. The amount of cells per microcarrier over time for various values for the proliferation threshold, similar to Fig 6 but with (A) n=1 and (B) n=2

**S4 Fig.**
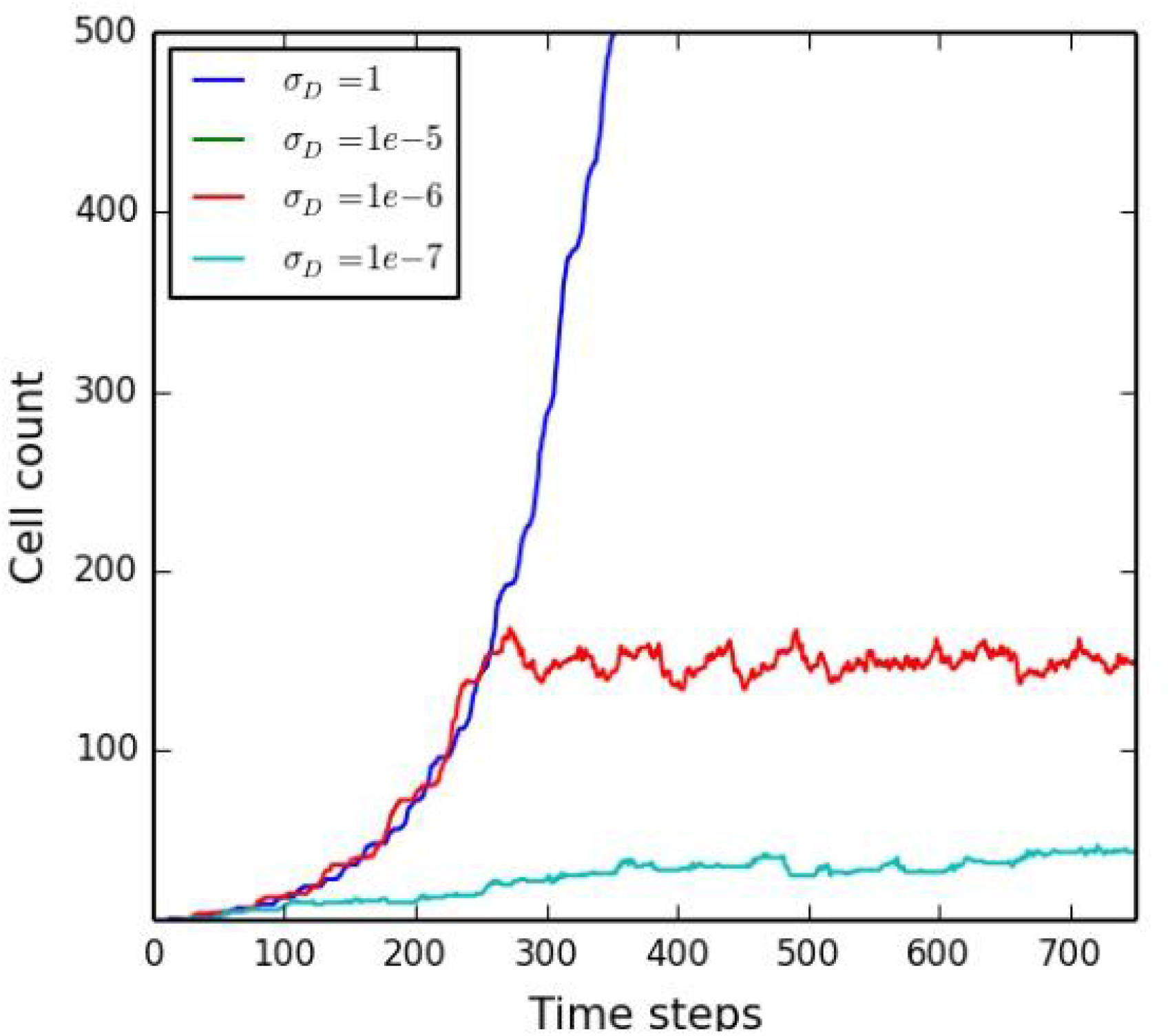
Cell population as a function of time for the cell death only scenario. The amount of cells per microcarrier over time for various values of σ_*D*_ and *T*_*D*_ = 0. 005.

**S5 Fig.**
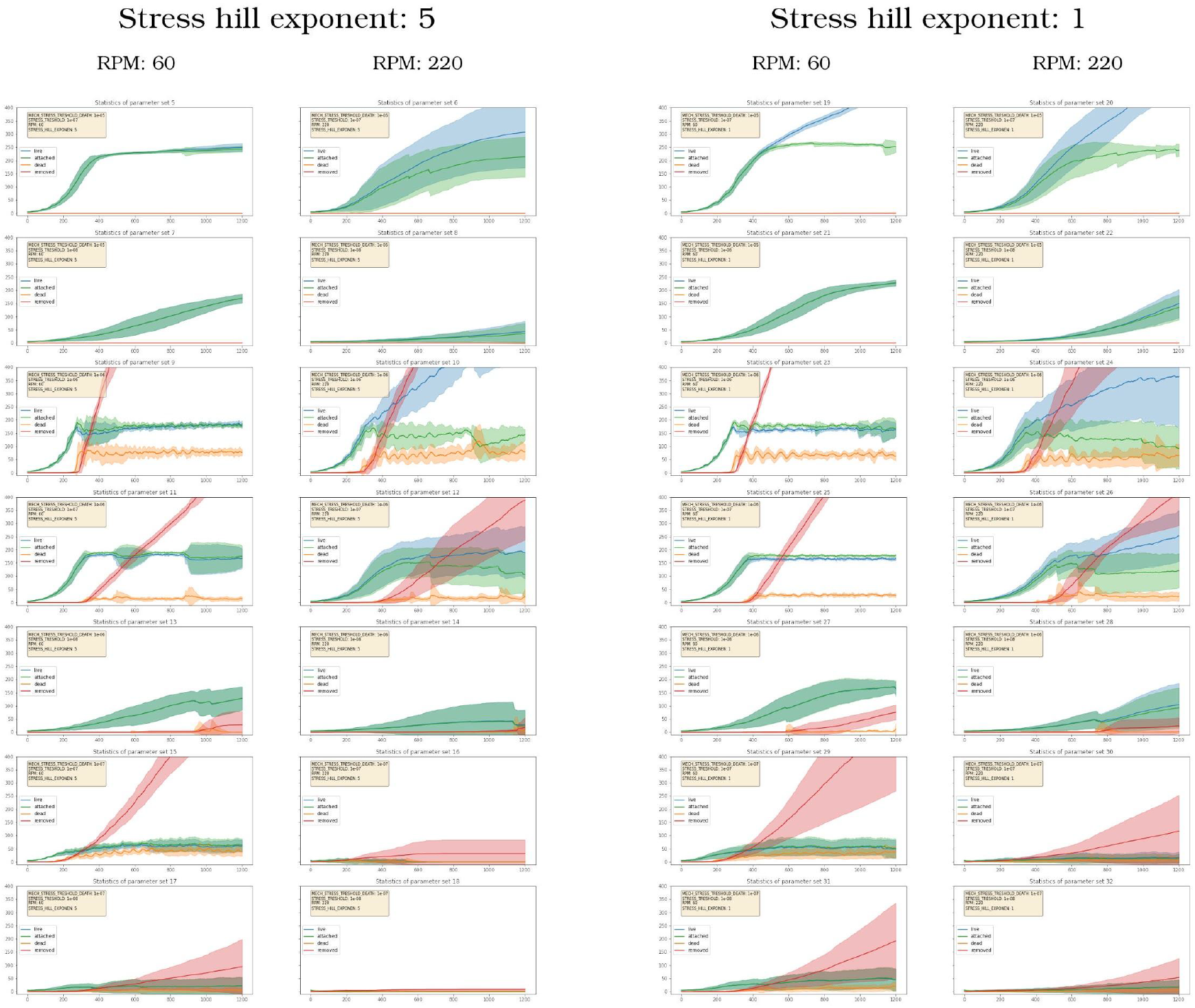
Cell population as a function of time for two rotors speeds. Simulations for two values, 5 and 1, of the stress hill component (that governs the steepness of the cell reaction profile to stress), two different rotor speeds and various other parameters as given in each yellow box. The solid line represents the mean value over all cells in 10 simulations, while the shaded band depicts the standard deviation from this mean. Cell counts are shown for the full simulation time for cells that are alive (blue), alive and attached to a microcarrier (green), undergoing apoptosis (orange) and cells that have died and are removed from the simulation (red).

**S6 Table.** Parameters and value ranges taken into account for the sensitivity analysis.

| Description (S2 Table) | Parameter name in source code | range_min | range_max | range_type |
| --- | --- | --- | --- | --- |
|  | STRESS_HILL_EXPONENT | 1 | 10 | linear |
|  | MECH_STRESS_TRESHOLD_DEATH | 1.00E-07 | 1.00E-05 | exponential |
| Mechanical stress modulator | STRESS_TRESHOLD | 1.00E-08 | 1.00E-06 | exponential |
| Cell-cell spring constant | A_AGENT_STIFFNESS_CC | 1.00E-04 | 1.00E-02 | exponential |
| Cell-cell bond flexibility | A_AGENT_BOND_S_CC | 1.00E-02 | 9.00E-01 | exponential |
| Thresholding factor for cell-cell bond breaking | A_AGENT_BOND_DESTROY_FACTOR_C<br>C | 1.00E+00 | 1.50E+00 | linear |
| Cell-microcarrier spring constant | A_AGENT_STIFFNESS_CM | 2.20E-04 | 2.20E-02 | exponential |
| Cell-microcarrier bond flexibility | A_AGENT_BOND_S_CM | 1.00E-02 | 9.00E-01 | exponential |
| Thresholding factor for cell-microcarrier bond breaking | A_AGENT_BOND_DESTROY_FACTOR_C<br>M | 1.00E+00 | 1.50E+00 | linear |

**S7 Text. Software management and reproducibility**.

The CFD model was solved by a commercial finite element method (FEM)-based software COMSOL Multiphysics (v5.5). The geometry domain for the bioreactor was discretized into tetrahedral-shaped elements forming a mesh. A total of ∼1.7 million elements were generated. The model was performed on a computer workstation with a 128 GB RAM operating memory running on two 16-core Intel(R) Xeon(R) 3.20 GHz.

The ABM model was built using the Biocellion (v3.1) agent based modeling framework (https://biocellion.com/). The models and supporting scripts to run parameter sweeps and plot the figures can be found on GitHub:

- https://github.com/TheCMMC/ABM-only-microcarriers
- https://github.com/TheCMMC/ABM-microcarriers
- https://github.com/TheCMMC/biocellion-tools

